# Mitochondria serve as axonal shuttle for Cox7c mRNA through mechanism that involves its mitochondrial targeting signal

**DOI:** 10.1101/2021.05.19.444640

**Authors:** Bar Cohen, Adi Golani-Armon, Topaz Altman, Anca F. Savulescu, Musa M. Mhlanga, Eran Perlson, Yoav S. Arava

## Abstract

Localized protein synthesis plays a key role in spatiotemporal regulation of the cellular proteome. Neurons, which extend axons over long distances, heavily depend on this process. However, the mechanisms by which axonal mRNAs are transported to protein target sites are not fully understood. Here, we describe a novel role for mitochondria in shuttling a nuclear encoded mRNA along axons. Fractionation analysis and smFISH revealed that the mRNA encoding Cox7c protein is preferentially associated with mitochondria from a neuronal cell line and from primary motor neuron axons. Live cell imaging of MS2-tagged Cox7c or Cryab control mRNA in primary motor neurons further confirmed the preferential colocalization of Cox7c mRNA with mitochondria. More importantly, Cox7c demonstrated substantial cotransport with mitochondria along axons. Intriguingly, the coding region, rather than the 3’UTR, was found to be the key domain for the cotransport. Furthermore, we show that puromycin treatment as well as hindering the synthesis of the mitochondrial targeting signal (MTS) reduced the colocalization. Overall, our results reveal a novel mRNA transport mode which exploits mitochondria as a shuttle and translation of the MTS as a recognition feature. Thus, mitochondria may play a role in spatial regulation of the axonal transcriptome and self-sustain their own proteome at distant neuronal sites.

## Introduction

Neurons extend axons over long distances and through a variety of extracellular microenvironments, forming unique compartmentalized sites with specific functions. These neuronal extensions rely on proper mitochondrial activity for their growth, function and survival. Indeed, mitochondrial dysfunction can lead to neurodegeneration and motor neuron diseases (Altman et al., 2019; Misgeld and Schwarz, 2017; Tan et al., 2014). In addition to addressing metabolic and energetic needs, mitochondria also serve as important signaling hubs, and are required for local protein synthesis events (Gershoni-Emek et al., 2018; Merrill and Strack, 2014; Rangaraju et al., 2019; Sheng, 2014; Spillane et al., 2013). However, it is largely unknown how mitochondria replenish their proteome or alter it in response to local needs. This question is particularly intriguing for distant mitochondria, since most of their proteins are derived from nucleus-transcribed mRNAs, yet protein and mitochondria transport times from the soma are prolonged (Maday et al., 2014; Terenzio et al., 2017; Twelvetrees et al., 2012). These mitochondria are likely replenished by local synthesis of proteins, since mRNAs encoding mitochondrial proteins were identified at distant axonal sites and local synthesis appears necessary for mitochondrial function (Aschrafi et al., 2008a; Aschrafi et al., 2016; Gumy et al., 2011; Hillefors et al., 2007; Kaplan et al., 2009; Minis et al., 2014; Rotem et al., 2017; Shigeoka et al., 2016,)Golani-Armon and Arava, 2016). An accepted model for transport of these mRNA to neuronal extensions involves the formation of RNA granules that shuttle through an interaction with kinesin motors (Kanai et al., 2004; Mofatteh and Bullock, 2017; Sahoo et al., 2018). Recently, an endosome-mediated mechanism for protein replenishment was suggested, in which mRNAs encoding mitochondrial proteins are transported to distant sites through their association with Rab7-containing endosomes. These mRNAs are then locally translated, and the protein is targeted to nearby mitochondria (Cioni et al., 2019).

Previous data demonstrating that mRNA are associated with mitochondria (Eliyahu et al., 2012; Lesnik et al., 2014; Marc et al., 2002; Vardi-Oknin and Arava, 2019; Williams et al., 2014), and that miRNAs are associated with mitochondria along the axons (Gershoni-Emek et al., 2018) led us to propose a mitochondria-centric mechanism that is independent of other organelles. Specifically, we hypothesized that axonal mitochondria are physically associated with mRNAs encoding mitochondrial proteins. Moreover, we reasoned that these may cotransport along the axon. Consistent with that, we found significant association of several mRNAs encoding mitochondrial proteins with mitochondria from a neuronal cell line and primary motor neurons. MS2-tagging and live imaging of Cox7c (Cytochrome c oxidase subunit 7C, a component of complex IV of the respiratory chain) mRNA revealed significant cotransport with mitochondria. Intriguingly, the coding region and in particular translation of the mitochondrial targeting signal (MTS), appeared key to mRNA localization and cotransport. Thus, Cox7c represents a novel mode of mRNA transport within axons.

## Results

### Mitochondrial association of mRNAs encoding mitochondria proteins

To examine the possibility that mitochondria serve as an mRNA transport vehicle, we first performed a biochemical analysis of mRNAs’ association with mitochondria. Briefly, N2a cells, a mouse neuroblastoma cell line, were fractionated by differential centrifugation to purify the mitochondria. The fraction purity was verified by western blot analysis for compartmental protein markers, revealing an enrichment of the mitochondrial protein ATP5A and the absence of the cytosolic marker GAPDH (Fig. 1A, B). Furthermore, RT-qPCR for mRNAs transcribed inside the mitochondria: Nd5 (encoding NADH Dehydrogenase, Subunit 5) and Cox1 (encoding Cytochrome c oxidase I) also revealed an enrichment of these mRNAs in the mitochondrial fraction (Fig. 1C). Next, we analyzed mRNAs that encode mitochondrial proteins and are transcribed in the nuclear genome. To this end, we selected mRNAs encoding components of the oxidative phosphorylation pathway; studies in yeast revealed that such mRNAs are enriched near mitochondria (Williams et al., 2014). Importantly, the selected mRNAs were previously found to be enriched in rodents’ axons [(Cox4i1 (Cytochrome c oxidase subunit 4 isoform 1), Atp5k (ATP synthase subunit e), Cox7c (Cytochrome c oxidase subunit 7C)] (Aschrafi et al., 2010; Aschrafi et al., 2016; Natera-Naranjo et al., 2012; Rotem et al., 2017). As shown in Fig. 1D, these mRNAs appeared to be enriched in the mitochondrial fraction. Interestingly, other mRNAs that were found before in axon but do not encode proteins that reside in mitochondria, such as Cryab protein (α Crystallin B chain), Rplp0 (Ribosomal protein of the large subunit), and β actin were detected at low levels in the mitochondrial fraction (Fig. 1D). Although the significance of the association of these “hitchhikers” remains to be determined, all of them appear at lower levels than mRNAs encoding *bona fide* mitochondrial protein.

**Figure 1:**
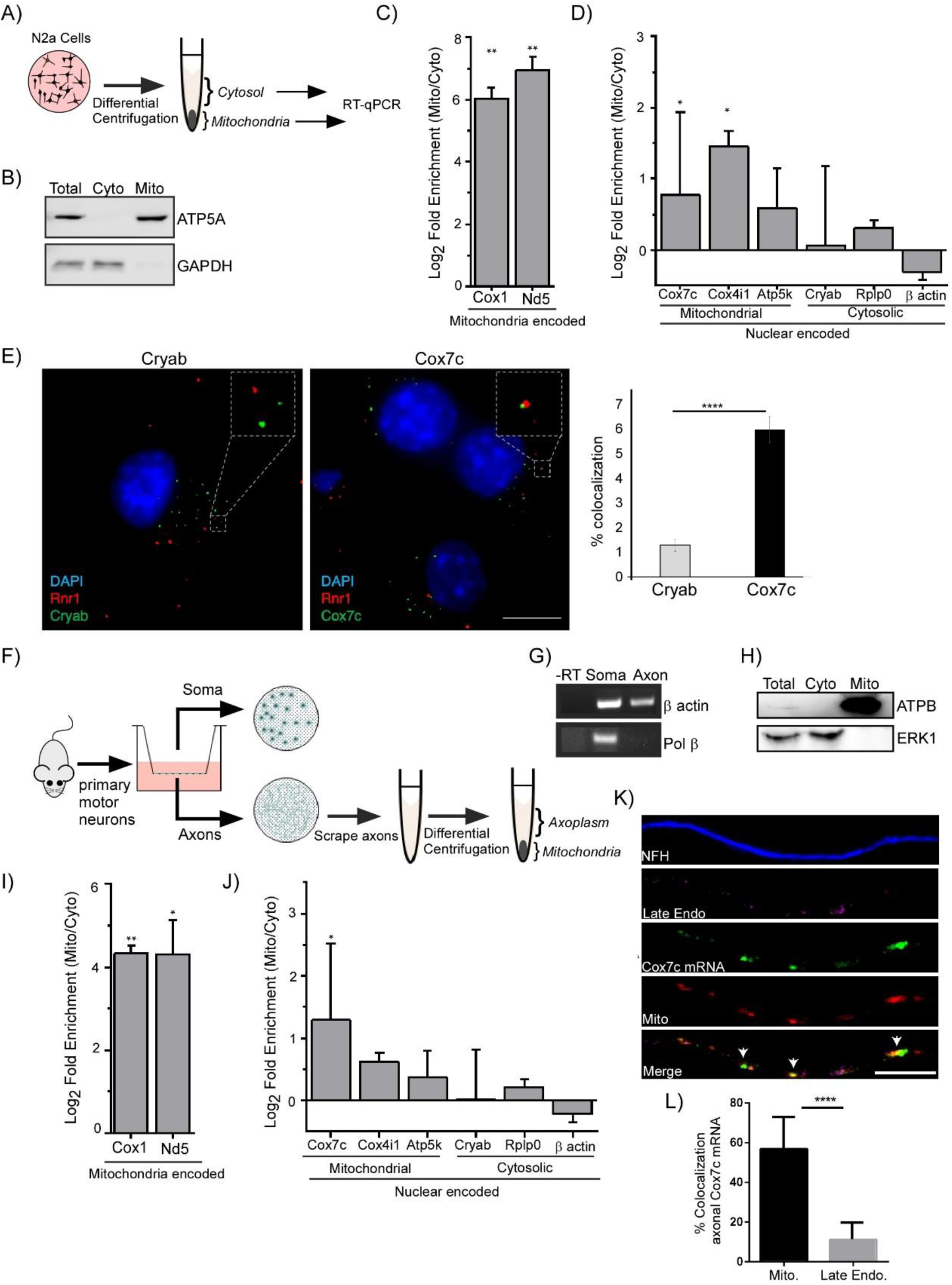
mRNAs encoding mitochondrial proteins are associated with mitochondria from N2a and motor neuron axons. A) Schematic workflow: N2a cells were collected and fractionated by differential centrifugation into cytosolic and mitochondrial fractions followed by RNA purification and RT-qPCR. B) Western-blot analysis of the unfractionated total sample, cytosolic (Cyto) and mitochondrial (Mito) fractions with a mitochondria marker (ATP5A) and cytosol marker (GAPDH), demonstrating fractionation purity. C) RT-qPCR analysis of mitochondrial and cytosolic fraction for mRNAs encoded inside the mitochondria. All values are normalized to β-actin and Rplp0 transcript levels and presented as log2 fold enrichment of mitochondria and cytosolic Ct signals. Error bars are SEM. Two-way ANOVA, \**p*<0.05, \*\**p*<0.01. n = 4 independent biological repeats. D) RT-qPCR analysis of mitochondrial and cytosolic fraction for mRNAs encoded in the nucleus, separated to those that their proteins are destined to mitochondria or cytosol. Quantification as in panel C. Error bars are SEM. Two-way ANOVA, \**p*<0.05. n = 4 independent biological repeats. E) Representative images of smFISH done on N2a cells with the mRNA (green) of Cryab (left) or Cox7c (right). Probes targeting the 21S mitochondrial rRNA (Rnr1-red) were used to detect mitochondria signal. The histogram presents the percent of spots of each mRNA that colocalize with Rnr1 signals. Student’s t-test, \*\*\*\**p*<0.0001. For Cryab, 2027 spots from 82 cells were analyzed, and for Cox7c 1747 spots from 69 cells were analyzed, in three independent experiments. To minimize ambiguity, only spots with overlapping signals were considered as colocalized. Scale bar = 5 μm. F) Schematic workflow used for membrane-based compartmental isolation of primary motor neuron axons, and their fractionation into axoplasm (Cyto) and mitochondrial (Mito) fractions followed by RNA purification and RT-qPCR. G) RT-PCR analysis for soma-specific mRNA (Pol β) confirming separation purity. β-actin represents mRNA present in both fractions. H) Western-blot analysis for fractionated axonal samples, with a mitochondrial marker (ATPB) and cytosolic marker (ERK1) demonstrating purity of mitochondria samples. I) RT-qPCR analysis was performed on axonal RNA samples from mitochondrial and axoplasm fractions for mRNAs encoded in the mitochondria. All values are normalized to β-actin and Rplp0 transcript levels. Error bars are SEM. Two-way ANOVA, \**p*<0.05. n = 3 independent biological repeats. J) RT-qPCR analysis of mitochondrial and cytosolic fractions for mRNAs encoded in the nucleus, separated to those that their proteins are destined to mitochondria or cytosol. Quantification as in panel I. Error bars are SEM. Two-way ANOVA, \**p*<0.05. n = 3 independent biological repeats. K) Representative images of smFISH done on primary motor neurons for Cox7c mRNA (green) along with immunostaining for axons by neurofilament heavy chain (NFH, blue), late endosomes (Rab7 marker, magenta) and mitochondria (MitoTracker staining, red). Arrows indicate areas of colocalization between mitochondria and Cox7c mRNA. Scale bar =10µm. L) Colocalization analysis of Cox7c mRNA with mitochondria and late endosomes signals. *n=*41 axons from 3 repeats. Error bars are SD. Unpaired-*t-test ****p<0*.*0001*.

To validate the biochemical association data by an alternative approach, we performed smFISH to visualize Cox7c, a representative of mRNAs encoding proteins that are inserted into the mitochondria, and Cryab, a control mRNA. The colocalization of each of these transcripts with RNA transcribed by the mitochondrial genome (Rnr1) that served as a mitochondrial marker was quantified; The smFISH data revealed a higher colocalization for Cox7c than for Cryab (Fig. 1E), consistent with the fractionation results.

Next, we investigated whether this association occurs in mitochondria from axons of mouse motor neurons. To that end, we developed a protocol for isolating mitochondria with their associated mRNAs from primary spinal cord axons. Cells were extracted from embryonic spinal cords and grown on a porous membrane (“Modified Boyden Chamber”) that enabled us to separate the axons from the cell bodies. Soma and axonal protrusions were then scraped (Fig. 1F) (Gershoni-Emek et al., 2018; Rotem et al., 2017). RT-PCR analysis confirmed the absence of soma-marker mRNA (Pol β) from the axonal fractions; however, a known axonal mRNA (β actin) was clearly detected in axonal fractions (Fig. 1G). A fractionation protocol was devised to fractionate axons to mitochondria and axoplasm; western analysis confirmed the absence of a cytosolic marker from the mitochondrial fraction and vice versa (Fig. 1H) and RT-qPCR revealed a significant enrichment for mRNAs encoded by the mitochondrial genome in the mitochondrial fraction (Fig. 1I). Importantly, although variation between biological repeats is apparent due to complexity of the protocol, axonal mRNAs encoding mitochondrial proteins (predominantly Cox7c and Cox4i1) appear enriched in the mitochondrial fraction, whereas those encoding non-mitochondrial proteins (Cryab, Rplp0, and β actin) are at similar levels in the mitochondrial and axoplasm fractions (Fig. 1J). Thus, mRNAs encoding mitochondrial proteins are associated with mitochondria from primary neurons axons.

Recently, it was shown that mRNAs encoding mitochondrial proteins, such as *vdac2*, are bound to late endosomes in neuronal processes (Cioni et al., 2019). To assess if Cox7c mRNA possesses similar qualities, we cultured motor neurons and performed smFISH of Cox7c mRNA together with mitochondrial staining, and immunostaining for the late endosomal marker Rab7. Indeed, a fraction of Cox7c appeared to be colocalized with late endosomes. However, a much higher fraction appeared to colocalize with mitochondria (Fig. 1K-L). Thus, Cox7c mRNA is bound to both organelles and might be transported in axons by an alternative pathway, independent of endosomal mediated transport.

### mRNAs cotravel with mitochondria along axons

To explore cotransport of mRNAs with mitochondria, we utilized the MS2 system (Wu et al., 2012) to tag candidate mRNAs and follow their position and transport by confocal live imaging. The complete coding region (CDS) of Cox7c and Cox4i1 (as representatives of highly associated candidates) and of Cryab (as a low association, control marker) were cloned in-frame to CFP in an MS2-loop bearing expression vector (Fig. 2A). Furthermore, the 3’UTR of each gene was cloned downstream to the MS2-loops. This yielded transcripts (designated ‘Full’) that contain the entire transcript of each gene tagged with multiple MS2 loops. Each construct was transfected into N2a cells together with a plasmid expressing MS2-binding protein fused to YFP (MCP-YFP). Confocal microscopy was used to measure colocalization with MitoTracker-red stained mitochondria (Fig. 2A). Quantification of the mRNA signals that colocalized with mitochondrial signals revealed about 25 % colocalization for the control, parental vector that expresses only the MS2 loops (Fig. 2B). This level is most likely due to the abundance of mitochondria and MCP-YFP and represents the basal background of the system. Nevertheless, the MS2 constructs expressing Cox7c or Cox4i1 exhibited significantly higher colocalization values. Colocalization of Cryab mRNA was non-significantly different from the control MS2-only mRNA. Overall, these data support the preferred association of nuclear encoded mitochondrial mRNAs with mitochondria in a neuronal cell line.

**Figure 2:**
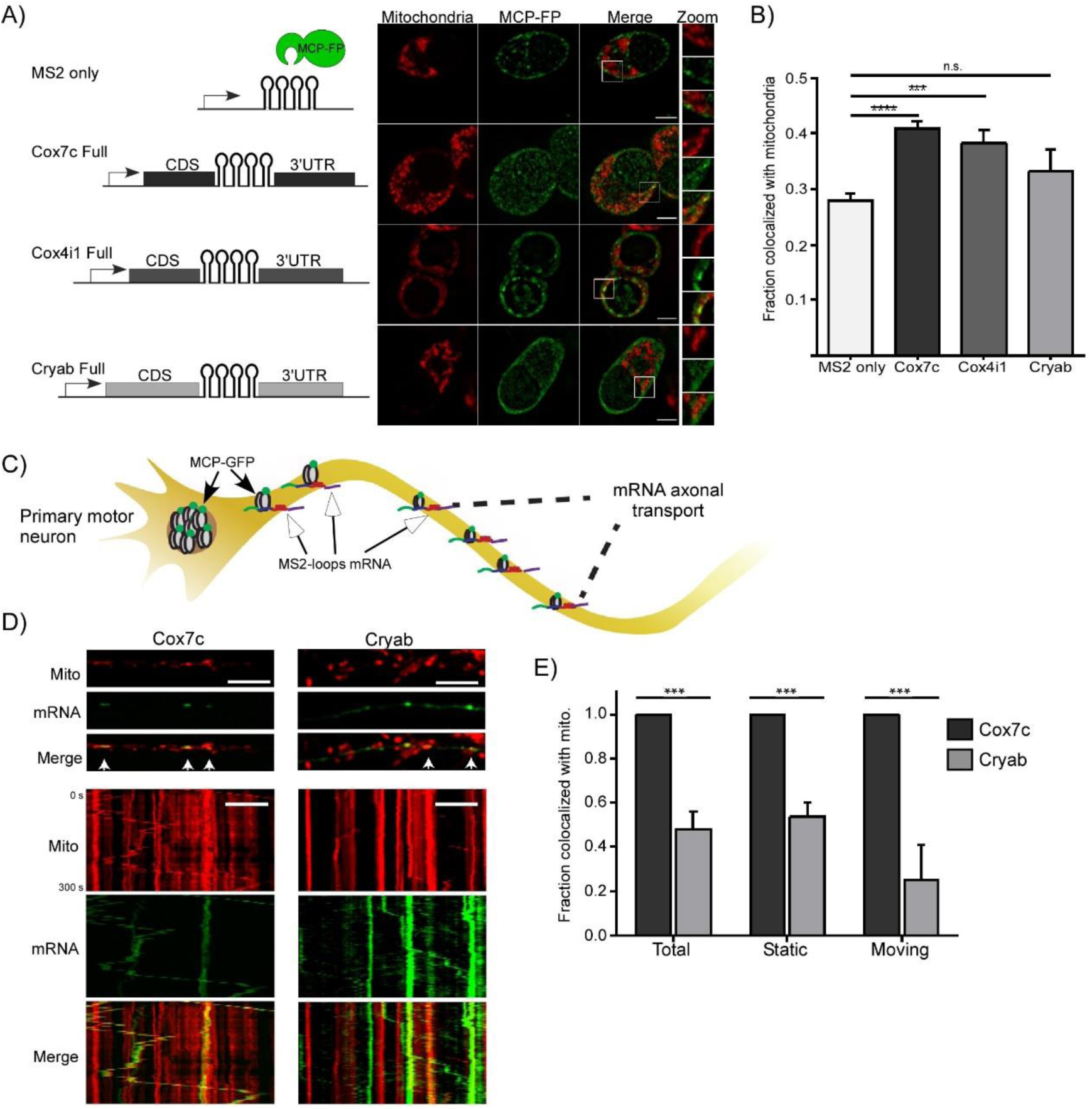
Live imaging reveals cotransport of mRNA with mitochondria. A) Left: Schematic representation of the genetic constructs encoding the coding sequence (CDS) and the 3’UTR of Cox7c, Cox4i and Cryab genes, which were cloned with multiple MS2 aptamers. Right: Representative images of the indicated MS2-mRNA constructs expressed in N2a cells together with a vector expressing MS2 binding protein fused to YFP (MCP-FP – green) colocalized with mitochondria staining (red). Scale bar = 5 μm. B) Quantification of the mRNA signal overlapping with mitochondria. Error bars are SEM. Mann-Whitney test, n.s (non-significant) *p*>0.05, \*\*\**p*<0.001, \*\*\*\**p*<0.0001, n = 34, 48, 9, 14 cells for MS2 only, Cox7c, Cox4i1 and Cryab, respectively. C) Scheme of mRNA imaging in primary motor neurons. MS2-loops containing constructs are introduced by viral infection into primary motor neurons together with a plasmid expressing GFP-tagged MS2 binding protein (MCP-GFP). Cells are grown in microfluidic chambers and signals are quantified along axonal extensions by spinning-disc microscopy. D) Representative images and kymographs of primary motor neurons infected with viral vectors expressing MCP-GFP and Cox7c or Cryab MS2-mRNA constructs (green), showing colocalization with axonal mitochondria (red). Arrowheads indicate colocalization of mRNA and mitochondria. Scale bar = 10 μm. E) Quantification of axonal cotransport between mRNA and the mitochondria separated to moving and static mRNA. Error bars are SEM. Two-way ANOVA, \*\*\**p*<0.001. n = 7 and 9 axons for Cox7c and Cryab, respectively, from 3 biological independent cultures, and > 100 MS2-mRNA particles for each construct.

We next introduced the plasmids with MS2-tagged genes and MCP-GFP by viral infection into primary spinal cord neurons (Fig. 2C). Live imaging using spinning disk confocal microscopy provided high spatial and temporal resolution of mitochondria and subcellular localization of tagged mRNAs. We selected Cox7c for detailed analysis since it showed the highest enrichment among all tested mRNAs, yet its localization mechanisms are largely unknown (compared to Cox4i1 that has extensively been studied (Aschrafi et al., 2008b; Aschrafi et al., 2010; Kar et al., 2014)). Cryab was used as a non-mitochondrial control mRNA. Notably, all Cox7c mRNA signals appeared to colocalize with mitochondria, while less than 50 % of Cryab signals were colocalized. Furthermore, all moving mRNA signals of Cox7c were in complete concordance to mitochondria movement, while only a small fraction of Cryab was in concordance with mitochondria (Fig. 2D, E and Suppl. Movie 1,2). Thus, mRNA encoding mitochondria proteins not only associate with axonal mitochondria, but also appear to shuttle with them along the axon.

### The coding region contains key cotransport determinants

To identify which mRNA regions are important for cotransport of Cox7c with mitochondria, either the Cox7c coding region (CDS) alone or the 3’UTR alone were cloned into an MS2-loop bearing expression vector and expressed in N2a cells. Northern blot analysis confirmed that all transcripts (Full, CDS only or 3’UTR only) are of the expected length (data not shown). To determine the localization of these mRNA variants, N2a cells were transfected with these constructs and with the MCP-YFP plasmid. Live imaging revealed that whereas the mRNA carrying the CDS region exhibited similar colocalization values to those of the entire Cox7c mRNA (‘Full’), the mRNA carrying only the 3’UTR region had a significantly lower association (Fig. 3A, B). This indicates that the coding region is sufficient to induce mitochondrial localization. As a complementary approach to imaging, we biochemically fractionated the transfected cells into cytosolic and mitochondrial fractions. Fractions were subjected to western blot analysis for MCP-YFP, as a proxy for the amounts of mRNA in each fraction (Fig. 3 C, D). The CDS-only and the Full transcripts were higher than the 3’UTR only (or the MS2 only control) in the mitochondrial fraction. This fractionation data further supports the live imaging data indicating a role for the coding region in mRNA localization to mitochondria.

**Figure 3:**
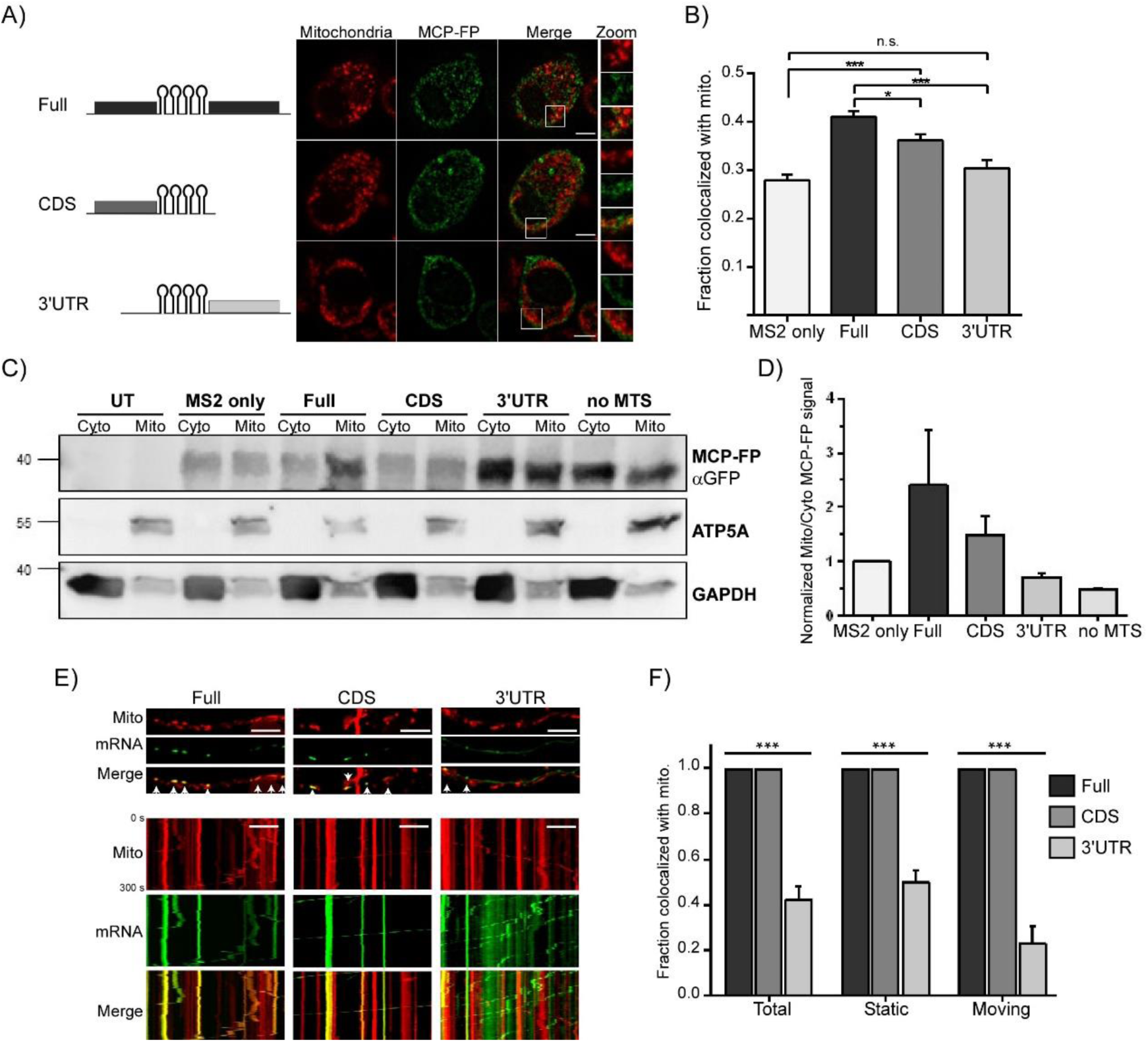
The coding region (CDS) contains colocalization determinants. A) Left: Schematic representation of MS2-loops Full Cox7c, Cox7c CDS only, or Cox7c 3’UTR only constructs. Right: Representative images of N2a cells transfected with the MS2 constructs. Scale bar = 5 μm. B) Quantification of the mRNA signal overlapping with mitochondria. Error bars are SEM. Mann-Whitney test, n.s. (non-significant) p>0.05, \**p*<0.05, \*\*\**p*<0.001, n = 23, 48, 17, 25 cells for MS2 only, Cox7c, CDS and 3’UTR, respectively. C) Western blot analysis of cytosolic (Cyto) and mitochondrial (Mito) fractions. Mitochondria marker (ATP5A) and cytosol marker (GAPDH), demonstrating fractionation sufficiency. GFP antibody is used to identify the MCP-YFP in the different fractions (as a proxy for MS2-tagged mRNA localization). D) Quantification of MCP-FP signal, in cytosolic (Cyto) and mitochondrial (Mito) fractions. The signals were corrected to fractionation efficiency protein markers (cytosolic contamination in mitochondria fraction), and then ratios were normalized to MS2 ratio. Error bars are SEM. n = 2 independent biological repeats. E) Representative images and kymographs of primary motor neurons infected with viral vectors expressing either Full Cox7c, Cox7c CDS only, or Cox7c 3’UTR only and MCP-GFP (green). Axons show axonal colocalization with mitochondria (red) in the Full and CDS only but not in the 3’UTR only construct. Arrowheads indicate colocalization of mitochondria and mRNA signals. Scale bar = 10 μm. F) Quantification of axonal cotransport of mRNA and mitochondria, separated to moving and static mRNA. Error bars are SEM. Two-way ANOVA, \*\*\**p*<0.001. n = 8, 12, 8 axons from 3 biological independent cultures, for Cox7c Full (95 particles), Cox7c CDS (215 particles) and Cox7c 3’UTR (115 particles), respectively.

Importantly, we introduced the deletion variants into primary motor neurons (Fig. 3E, F, Supplementary movies 3 and 4). Also here, the CDS-only colocalization signals appeared very similar to those of the Full construct. Furthermore, focusing on the motile mRNAs in axons revealed that all these transcripts are moving together with mitochondria. However, this was not the case for the 3’UTR variant: most of its signal did not correlate with the movement of mitochondria. Overall, these data reveal that elements within the coding region are important for cotransport of Cox7c transcript with mitochondria.

### Colocalization is dependent on translation of the Mitochondrial Targeting Signal (MTS)

The effects imposed by the CDS region suggest that translation of protein sequences contribute to colocalization. We therefore tested the possible role of a translational process by treating cells with the translation inhibitor puromycin, which leads to dissociation of ribosomes from mRNAs. While no effect was apparent in the MS2-only control, a significant reduction in mitochondrial association was observed for the mRNA expressing the complete Cox7c transcript (Fig. 4A, B).

**Figure 4:**
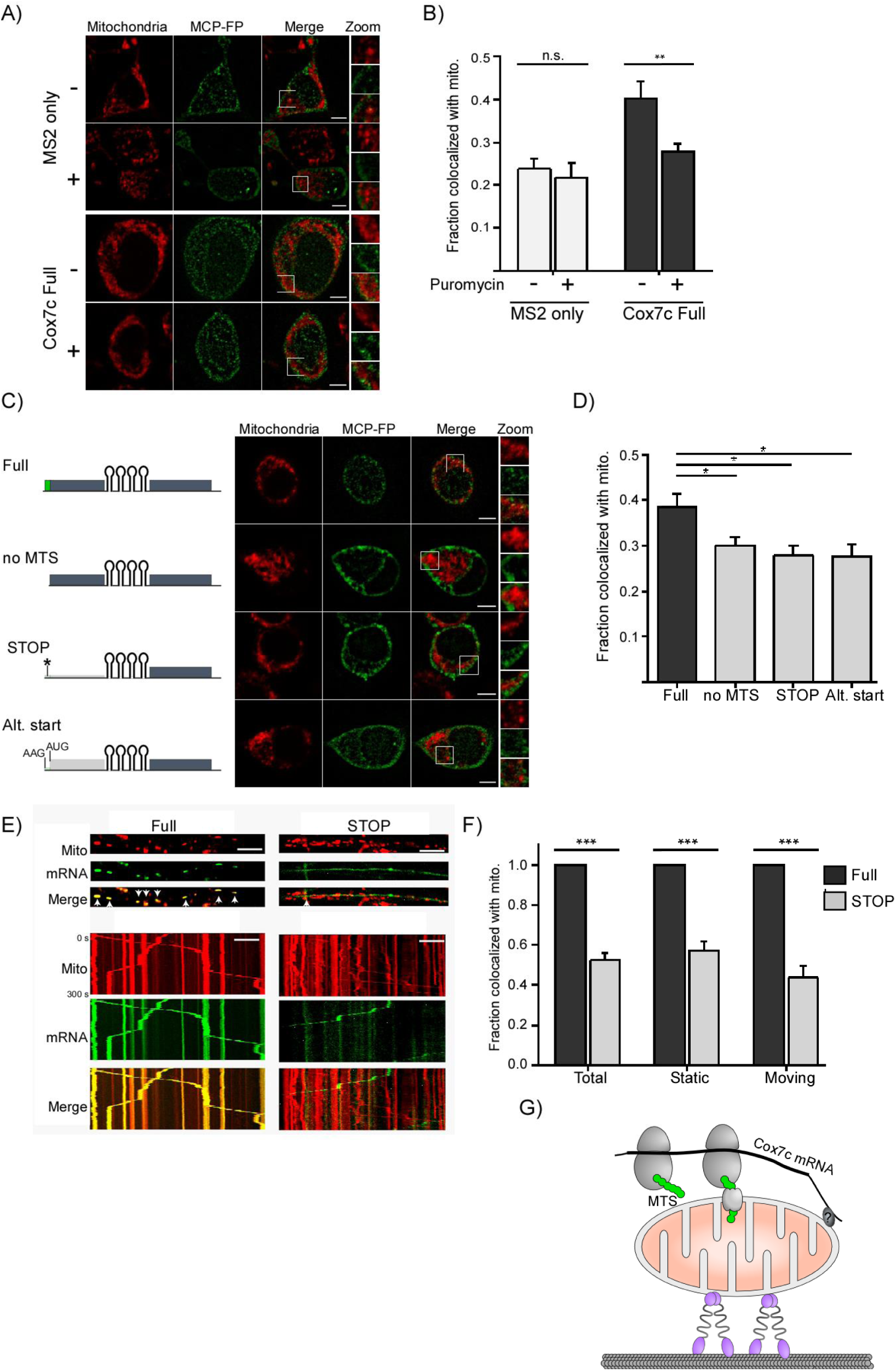
Translation of the MTS is important for localization. A, B) Cells transfected with the indicated plasmids were treated with the translation inhibitor puromycin (+ or -), and MCP-FP signals colocalized with mitochondria were quantified. Error bars are SEM. Mann-Whitney test, n.s (non-significant) *p*>0.05, \*\**p*<0.01, n = 12, 13, 17, 19 cells for MS2-only without or with puromycin and Cox7c without or with puromycin, respectively. C) Left: Schematics of the Cox7c MTS variants used: ‘no MTS’ - without 15 amino acids predicted to serve as MTS; ‘STOP’ – a point mutation generating a stop codon closely downstream to the initiation codon; ‘Alternative Start’ - point mutations that shift the start codon downstream of the MTS. Right: Representative images of colocalization with mitochondria. Scale bar = 5 μm. D) Quantification of colocalization from 3 independent biological repeats. Error bars are SEM. Mann-Whitney test * *p* <0.05. n = 14, 11, 18, 13 cells for Cox7 Full, no MTS, STOP and Alt. start, respectively. E) Representative images and kymographs of primary motor neurons infected with viral vectors expressing Full Cox7c or Cox7c ‘STOP’ and MCP-GFP (green). Axons show axonal colocalization with mitochondria (red) in the Full Cox7c but not in the Cox7c ‘STOP’ construct. Arrowheads indicate colocalization of mitochondria and mRNA signals. Scale bar = 10 μm. F) Quantification of axonal cotransport of mRNA and mitochondria, separated to moving and static mRNA. Error bars are SEM. Two-way ANOVA, \*\*\**p*<0.001. n = 9 and 19 axons from 3 biological independent cultures, for Cox7c Full (170 MS2-mRNA particles) and Cox7c ‘STOP’ (231 particles), respectively. G) Suggested model for cotranslation-mediated mRNA transport with mitochondria. Translation of the nascent MTS (green) targets Cox7c mRNA to mitochondria and contribute to its cotransport with the organelle. Additional RNA binding proteins (denoted by ‘?’) are likely to anchor the mRNA to the outer membrane.

Most mitochondria proteins are targeted to the mitochondria through an N-terminal Mitochondria Targeting Signal (MTS) (Harbauer et al., 2014). Previous studies in yeast revealed the importance of MTS translation for directing the mRNA to mitochondria proximity (Eliyahu et al., 2010; Lesnik et al., 2014). To assess whether this mechanism underlies mRNA association with axonal mitochondria, we generated a Cox7c variant with a deletion of the 15 amino acids that are predicted to serve as an N-terminal MTS (Armenteros et al., 2019) (‘no MTS’). Confocal imaging of mRNA localization in N2a cells revealed a significant reduction in colocalization with mitochondria compared with the full, MTS-containing mRNA (Fig. 4C). Consistently, fractionation analysis of MCP-YFP, as a proxy for the mRNA localization, revealed significantly lower localization to the ‘no MTS’ compared to the Full construct (Fig. 3C, D). To exclude the possibility that the deleted 45 nts mRNA sequence underlies targeting, we generated a Cox7c variant with a non-sense point mutation at the fifth codon of Cox7c (‘STOP’), which leads to a premature translation termination but without any additional change in the Cox7c mRNA sequence. Furthermore, since this STOP mutation abolishes translation of the entire Cox7c protein, we generated a construct with an alternative start codon downstream to the MTS (Alt. Start), i.e., a missense mutation at the start codon (AUG to AAG) and a start codon at the 16^th^ codon were introduced. This leads to translation of Cox7c yet without its MTS. As seen in Fig. 4D, these two variants, which are almost identical to the natural Cox7c mRNA, show a clear decrease in mRNA colocalization. Thus, translation of the MTS itself, rather than the mRNA sequence or other Cox7c protein domain, directs Cox7c mRNA to mitochondria proximity.

Finally, introducing the ‘STOP’ construct to primary motor neurons revealed similar findings, i.e., reduced colocalization of mRNA particles with mitochondria (Figure 4E-F (‘Static’ bars), Supplementary movie 5). Moreover, axonal cotransport of this mRNA is also reduced compared to the full Cox7c construct (Figure 4E-F (‘Moving’ bars)). Thus, translation of the MTS may play a role in mitochondria localization and (consequently) cotransport in primary motor neurons.

## Discussion

The data presented here reveal a novel mode of mRNA transport in neurons. We propose a model (Fig. 4G) in which nuclear-encoded mitochondrial mRNA is associated and transported with axonal mitochondria. We describe new determinants, namely translation and the emerging MTS, for proper mRNA association and hence cotransport. Translation of a signal peptide is the main mechanism for mRNA targeting to the ER (Aviram and Schuldiner, 2017), and we previously reported a similar colocalization mode for mRNAs encoding mitochondrial proteins with mitochondria of yeast cells (Eliyahu et al., 2010; Eliyahu et al., 2012; Lesnik et al., 2014). Yet, to the best of our knowledge, involvement of these determinants in long distance traffic has never been shown. It should be noted that MTS association with the mitochondria outer membrane is transient, as this moiety is quickly inserted into mitochondria matrix and usually cleaved off the protein (Harbauer et al., 2014). Thus, we propose that the MTS serves primarily homing purposes, and once the mRNA is in proximity to mitochondria additional mRNA regions are employed for long term anchoring. Such localization regions are usually located in 3’UTRs and interact with RNA binding proteins on the mitochondria outer membrane (Glock et al., 2017; Mofatteh and Bullock, 2017; Sahoo et al., 2018)(Schatton and Rugarli, 2018). Indeed, a cis element located within the 3’ UTR of another Cox-family member (CoxIV) was shown to mediate its axonal localization (Aschrafi et al., 2010). Similarly, in the present work, the mRNA containing only Cox7c 3’UTR was found to have some association with mitochondria, and disruption of the MTS did not fully abolish axonal cotransport of the Cox7c ‘no MTS’ construct (Fig. 4). Thus, multiple, not necessarily mutually exclusive, mRNA regions may ensure mRNA association with mitochondria for long periods of time.

Notably, Harbauer *et al*. had found that Pink1 mRNA is cotransported with mitochondria in neurons (Angelika B. Harbauer et al., 2021). Together with the data presented herein, this suggests a broad phenomenon, in which mitochondria are associated with various mRNAs encoding mitochondria proteins. Thereby axonal mitochondria may be self-sustained with respect to their protein content, which would allow immediate response to local needs as mitochondria meander along changing axonal microenvironments. Furthermore, mitochondria might serve as a shuttle vector for diverse types of mRNA to neuronal extremities, and by that, spatiotemporally regulate cellular proteome. Indeed, we find several mitochondria-unrelated mRNAs to have some association with mitochondria (“hitchhikers”). A general role for mitochondria in mRNA shuttling is yet to be determined by more sensitive, genome-wide approaches. Another open question at this stage is whether the interaction of mRNA with mitochondria occurs already at the cell body, and whether this association is maintained throughout the lifetime of the mRNA. While we did not observe any mRNA dissociations through the time course of our analysis, higher resolution temporal and spatial tools are needed to follow mRNA traffic along the entire neuron.

Recently, a study in retinal ganglion cells of *Xenopus* revealed an endosome-mediated mode of transport for two mRNAs (*lamnb* and *vdac2*) with roles in mitochondrial function (Cioni et al., 2019). In addition, a genome-wide analysis in the plant pathogen *U. maydis* suggested that many more mitochondrial mRNA are transported by endosomes (Olgeiser et al., 2019). Thus, multiple pathways may work in parallel to ensure sufficient protein content for mitochondria at sites distant from the cell body (Müntjes et al., 2021). Notably, lysosomes were found to efficiently shuttle a non-mitochondrial mRNA (β-actin) in axons (Liao et al., 2019). Thus, mRNA trafficking by moving organelles emerges as a general concept in the neuronal transport of diverse mRNA types.

## Author contributions

B.C., A.G.-A., T.A., E.P. and Y.S.A. designed the work, B.C, A.G.-A. and T.A. performed the experiments and analyzed the data, A.F.S. performed the smFISH experiments and analyses in N2a cells, M.M.M., E.P., and Y.S.A provided funding, facilities and guidance, E.P and Y.S.A. wrote the manuscript with input from coauthors.

## Acknowledgments

We thank Profs. Shai Berlin (Technion) and Yaron Shavtal (Bar Ilan University) for cells and plasmids, Prof. Florence Rage for her kind help with smFISH experiments, Prof. Ishi Talmon (Technion) for advice, and members of our labs for ideas and support. This work was funded by the ISF (grant 1096/13)(YSA), Adelis Brain Research-grant 2023478 (YSA) and ISF (grant 735/19) to (EP).

## Methods

### Animals

HB9::GFP (stock no. 005029) mice colony was originally obtained from Jackson Laboratories. The colony was maintained by breeding with ICR mice (Jackson Laboratories). Mice were genotyped by DNA extraction and PCR using a utilized kit. For experimental purposes, only WT embryos of pregnant mice (GFP negative) were taken for motor neuron cultures.

Animal experiments were performed under the supervision and approval of the Tel-Aviv University Committee for Animal Ethics.

### Cell growth, transfection and treatments

Mouse neuroblastoma cells (N2a), a kind gift from Prof. Shai Berlin, Technion, were cultured in DMEM supplemented with 10% Fetal Bovine Serum (FBS), 2% Penicillin/Streptomycin, and 2 mM L-glutamine, at 37 °C in 5% CO2 atmosphere. Cells were split every 2-3 days by trypsinization. For transfection experiments, cells were grown to 60% confluency and transfected using jetPRIME reagent (Polyplus, 11415) according to manufacturer’s instructions. Protein synthesis inhibition treatments were performed by incubating the cells with 200 μg/ml puromycin in growth medium for 45 min before imaging.

### Primary motor neuron culture

Motor neurons were cultured from E12.5 ICR mouse embryos as previously described (Gershoni-Emek et al., 2018). Briefly, spinal cords were dissected, incubated with trypsin for 10 min, and triturated. The neuron containing soup was collected through a cushion of BSA, and the pellet was resuspended and centrifuged through a 10.4% Optiprep gradient for 20 min at 760 g with the brake turned off. The cells were collected from the interphase, centrifuged through a cushion of BSA, counted and plated. Motor neurons were grown and maintained at 37°C 5% CO2 incubator, in complete neurobasal medium containing 2% B27, 2% horse serum, 25 μM beta-mercaptoethanol, 1% Penicillin-Streptomycin, and 1% Glutamax supplemented with 1 ng/ml GDNF, 0.5 ng/ml CNTF, and 1 ng/ml BDNF.

### Lentiviral production and infection

Lentiviral constructs were produced in HEK cells using a second-generation packaging system with Gag-Pol and VSVG helpers. HEK cells were grown on a 100-mm dish at high confluence, replaced to antibiotics free medium and transfected with 10 µg of MS2 vector, 10 µg of MCP-GFP and 20µg of helper plasmids. Plasmids were placed in a calcium phosphate transfection mix (25 mM HEPES, 5 mM KCl, 140 mM NaCl, and 0.75 mM Na2PO4 with 125 mM CaCl2) in 1 mL volume per plate. Medium was replaced 8 hours after transfection, and supernatants were harvested 2 days afterwards. Lentiviral particles were concentrated 10-fold by using the PEG virus precipitation kit and kept at −80°C until use. Viral infection was performed using 10–20 µl of lentivirus per microfluidic chamber.

### Mitochondria fractionation from N2a cells

24 hours post-transfection, N2a cells were detached and washed with ice-cold phosphate buffered serum (PBS), and pelleted by centrifugation (600g for 5 minutes at 4°C). Cell pellet was resuspended in HM (0.6 M mannitol, 50 mM Tris-HCl pH=7.4, 5 mM MgAc, 100 mM KCl, 1 mM DTT, 1 gr/L BSA, 200 μg/ml CHX, 1 mM PMSF, 1 μM Leupeptin, 1 μM Pepstatin, 0.3 μM Aprotinin and 0.04 U/μl RiboLock RNase Inhibitor (ThermoFisher)), and lysed by 15-20 strokes of a Dounce homogenizer. Lysate was centrifuged twice (1,000g for 10 minutes at 4°C) to pellet intact cells and nuclei, and supernatant (designated “Total”) was further centrifuged at 15,000g for 15 minutes at 4°C to obtain a cytosolic fraction (supernatant) and a mitochondrial fraction (pellet).

### Axonal harvesting and mitochondrial fractionation

Porous membranes (“Modified Boyden chamber”) with 1 μm pores were placed in a 6 well culture plate. The membranes were coated with poly-DL-ornithine (1.5 μg/mL in PBS) overnight at 37°C then with Laminin (3 μg/mL in DDW) for at least 2 hours at 37°C prior to motor neuron plating. Primary motor neurons were then plated on the membranes (10^6^ neurons per a membrane insert) and grown with complete neurobasal media. After 10 days in vitro (DIV), membranes were washed 3 times with warm PBS, followed by gentle scraping of the axonal (bottom) part of the membrane into a tube containing mitochondria Homogenization Buffer (HM) (0.6 M mannitol, 50 mM Tris-HCL pH 7.4, 5 mM MgAc, 100 mM KCl, 1 gr/L BSA, 200 μg/ml cycloheximide (CHX), 1 mM DDT, 1X Protease inhibitor cocktail (Roche) and 0.04 U/μl RiboLock RNase Inhibitor (ThermoFisher)). Four membranes were collected for each experimental repeat. Axons collected in HM were sub-fractionated to cytosolic (supernatant, Cyto) and mitochondrial (pellet, Mito) fractions by two cycles of 12,000 g centrifugation for 15 minutes at 4°C.

### Western blot analysis

Samples from N2a cells and primary motor neurons were lysed and fractionated in mitochondria Homogenization Buffer (or immediately lysed after axonal harvest for somatic fraction), and were boiled in mixed in 1:4 (v/v) 5× Laemmli sample buffer for 5 min at 95°C. Next, samples were loaded on polyacrylamide gel and blotted on nitrocellulose membrane. The membrane was blocked with 5% BSA (N2a samples) or 5% milk (primary motor neuron samples) in TBS-Tween for 1 hour. Membranes of N2a cells samples were incubated for 1 hour at room temperature (RT) with primary antibodies for ATP5A (1:1000, Abcam Ab119688) and GAPDH (1:2000, Abcam Ab181602). The membranes were next incubated with secondary antibodies against mouse (1:20000, Sigma A5906) or rabbit (1:20000, Sigma A9169) for 45 minutes at RT and were exposed to ECL imager after 5 min incubation with EZ-ECL reagent (Biological Industries). Membranes of primary motor neuron samples were incubated overnight at 4°C with primary antibodies for ERK1/2 (1:10000, Sigma Aldrich) and ATPB (1:1000, Abcam). The membranes were next incubated with secondary anti-HRP antibody (1:10000, Jackson Laboratories) for 2 hours at RT and were exposed to ECL imager after 5 min incubation with ECL reagent (Thermo Fisher).

### RT-qPCR

RNA was extracted from all samples (fractionated cells or axons) using Trizol reagent according to the manufacturer’s protocol. RNA was reverse transcribed using Maxima First Strand cDNA Synthesis kit (Thermo scientific). To verify fraction purity, cDNA was subjected to PCR amplification with β-actin and Polymerase β primers followed by gel electrophoresis. Quantification of target genes was performed by real-time qPCR using SYBR green (Thermo Scientific). Targets were normalized to β-actin and Rplp0 mRNA levels, and log2 fold-change between mitochondrial and cytosolic fractions was calculated (ΔΔCt). All real-time (RT) primers are listed in Table 1.

**Table 1:**
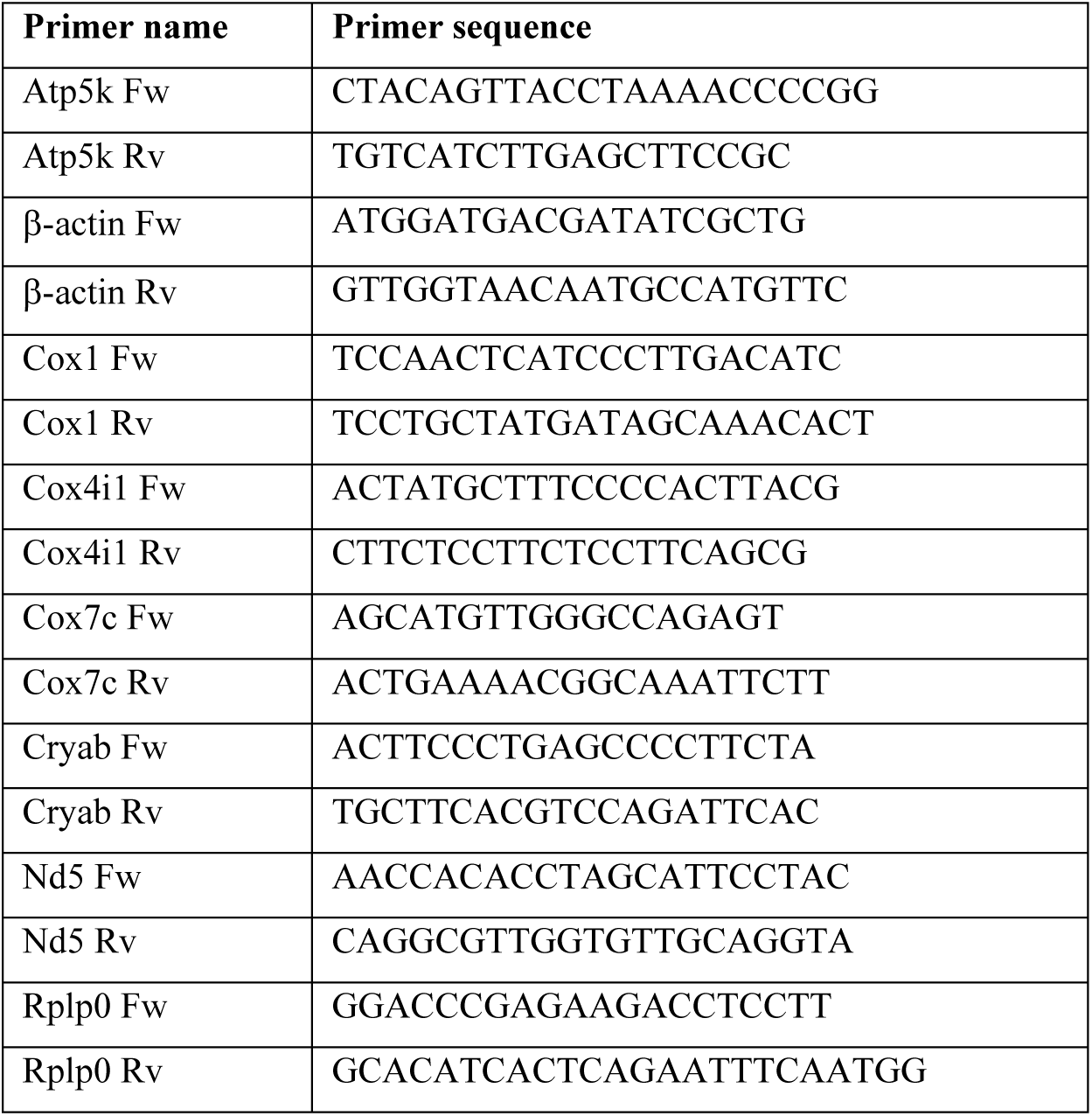
List of primers used for real-time PCR.

### Single molecule FISH (smFISH)

Design and manufacture of RNA FISH probes for use in the smFISH for N2a cells were performed according to the protocol by (Raj et al., 2008). Multiple 20-mer oligonucleotide probes conjugated to dyes, targeting the following mRNAs: Rnr1, Cryab and Cox7c were purchased (Biosearch Technologies). The Rnr1 probes were conjugated to TAMRA dye and the Cryab and Cox7c probes were conjugated to Quasar-670 dye. N2a cells were fixed in 3.7 % formaldehyde for 15 min at 37°C followed by washes in PBS and overnight permeabilization in 70 % Ethanol at 4°C. Cells were rehydrated in wash buffer (10 % formaldehyde, 2X SSC) for 5 min. Hybridization was conducted overnight in a humidified chamber at 37°C in Hybridization buffer (10 % dextran sulfate, 1 μg/μl E.coli tRNA, 2 mM Vanadyl ribonucleoside complex, 0.02 % RNase-free BSA, 10 % formamide, 2X SSC) combined with 50 ng of the desired RNA probe. Cells were then washed three times (each wash 30 min at room temperature) with FISH wash buffer (10 % formaldehyde, 2X SSC). Cells were then incubated in equilibration buffer (0.4 % glucose, 2X SSC) for 5 min and counter stained with 1 μg/ml DAPI (4’,6-diamidino-2-phenylindole; Life Technologies). Coverslips were mounted in imaging buffer (3.7 μg/μl glucose oxidase and 1U catalase in equilibration buffer) and imaged on a StellarVision microscope using Synthetic Aperture Optics (SAO).

Labeling of single mRNA molecules in mouse motor neurons was performed by smiFISH as previously described (Tsanov et al., 2016). Briefly, primary motor neurons were grown for 7 days on 13 mm coverslips. Labeling procedures were performed in RNase-free environment. Cultures were fixed with 4% PFA and permeabilized overnight with 70 % Ethanol. Samples were incubated with SSC (Sigma) based 15% Formamide (Thermo scientific) buffer for 15 minutes. Samples were hybridized overnight at 37°C with sixteen FLAP-Y-Cy-3 tagged complementary oligonucleotide probes, targeting regions in Cox7c mRNA (IDT) in Hybridization mix (15 % Formamide, 1.7 % tRNA (Sigma; R1753), 2 % FLAP:Probe mix, 1% VRC (Sigma), 1% BSA (Roche), 20% Dextran Sulfate (Sigma; D8906), 1x SSC). Samples were washed twice with warm 15% Formamide in 1xSSC buffer (1 hour each), then 30 minutes with 1x SSC buffer, and 30 minutes with 1x PBS. Prior to immunostaining samples were washed with Tris-HCl (pH 7.5), 0.15 M NaCl buffer, and then permeabilized with same buffer supplemented with 0.1 % Triton. Samples were blocked by 2% BSA (in the same buffer) for 30 minutes. Samples were incubated with primary antibodies for Chicken-anti-NFH (1:1000, Abcam) and Mouse-anti-Rab7 (1:200, abcam) in blocking buffer overnight at 4°C, and then with fluorescent secondary antibodies in blocking buffer for 2 hours at room temperature. Samples were mounted with Vectashield Antifade reagent.

### Plasmid construction

All mouse CDS and 3’UTR used were amplified from mouse cDNA using the primers indicated in Table 2. Fragments were cloned into phage-cmv-cfp-24xms2 vector (Addgene plasmid # 40651) created by Robert Singer lab. CDS fragments (from start codon to penultimate codon) were inserted in frame, upstream to CFP at the AgeI and NotI sites (under CMV promoter). 3’UTR fragments were inserted downstream to the MS2 stem loops cassette at the ClaI site. Constructs designated as ‘Full’ contain both the CDS and 3’UTR of the respective gene. ‘CDS’ or ‘3’UTR’ designations indicate constructs with only these regions of the gene (cloned at the above restriction sites) and the CFP reporter. Proper construction was confirmed by sequencing and proper transcripts’ length was validated by northern blot. We note that all constructs contained 12 MS loops instead of the expected 24, presumably due to recombination during cloning.

**Table 2:**
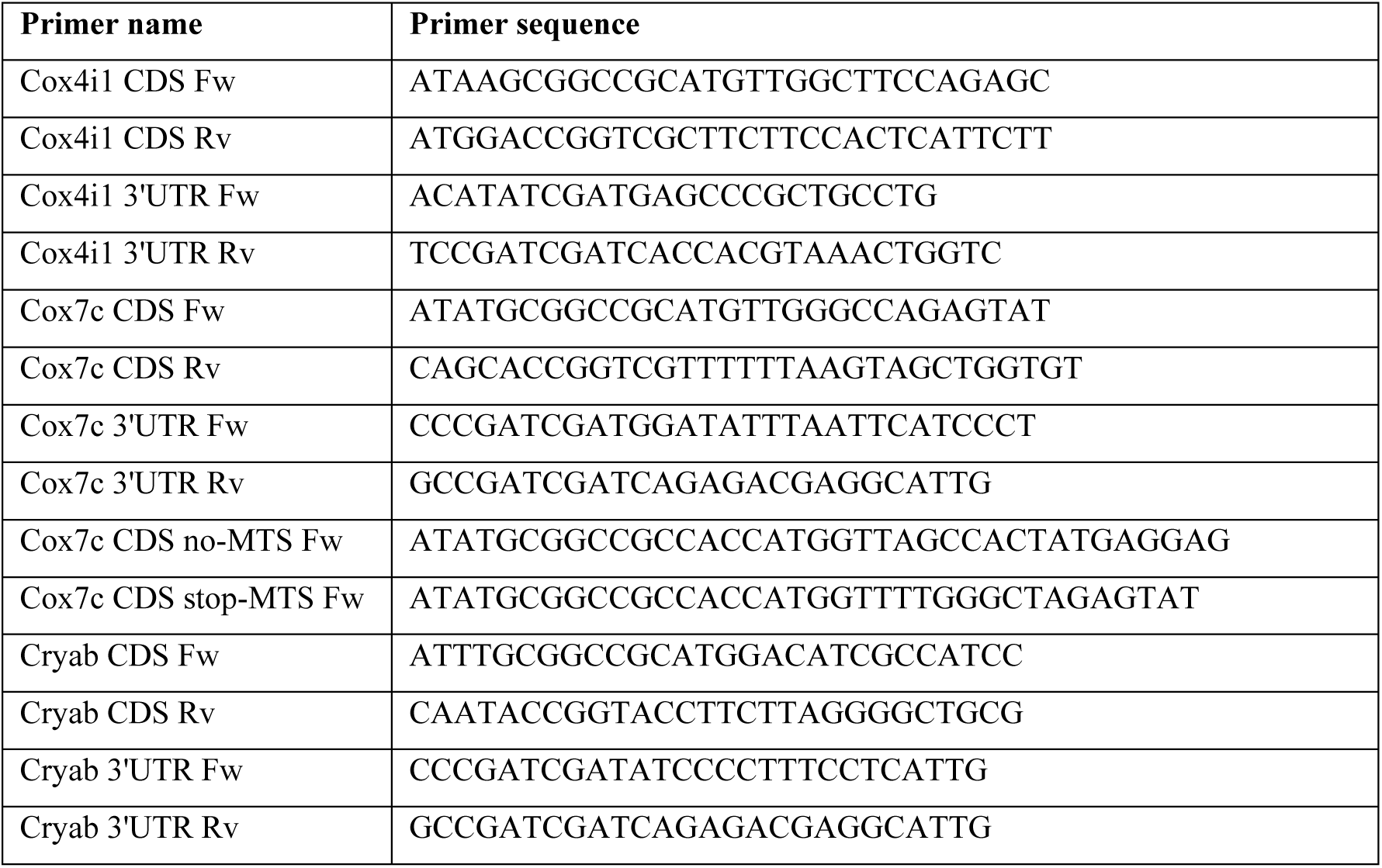
List of primers used for plasmid cloning.

MCP-YFP plasmid was a gift from Prof. Yaron Shav-Tal (Bar Ilan University). MCP plasmid used for axonal live-imaging was phage-ubc-nls-ha-tdMCP-gfp (Addgene plasmid # 40649).

### Motor neuron axonal transport in microfluidic chambers

Polydimethylsiloxane (PDMS) home-made microfluidic chambers (MFC) were casted as previously described (Ionescu et al., 2019). To enable motor neuron plating and medium exchange, four 6 mm wells were punched in the end of each channel surrounding the microfluidic grooves. The microfluidic devices were then cleaned, first with adhesive tape to remove dirt and then sterilized in 70% Ethanol for 10 min. The microfluidic devices were then dried, and UV irradiated for 10 min, followed by attachment to a 35 mm FluoroDish glass-bottom dish. For primary motor neuron plating, 150,000 neurons were concentrated and seeded in 4 µL medium to allow adhesion, and after 45 minutes 25 ng/mL BDNF enriched complete neurobasal medium was added. Lentiviral constructs were added to the proximal compartment two hours post-plating. From DIV 2, a neurotrophic and volume gradient was maintained between the distal and proximal compartment to encourage axonal crossing. Neurons were grown between 6-8 days to allow axonal crossing and lentiviral expression and were then stained with 100 nM MitoTracker Deep-Red FM for 30 minutes at 37°C followed by three washes with warm medium. For live axonal transport, movies of 100 frames were collected at 3 seconds intervals from the distal part of the MFC grooves. The movies were obtained using a Nikon Ti microscope equipped with a Yokogawa CSU X-1 spinning disc and an Andor iXon897 EMCCD camera controlled by Andor IQ3 software and 60x oil objective.

Axonal transport colocalization analysis was performed as previously described (Altman et al., 2019; Gershoni-Emek et al., 2018). Briefly, transport movies were analyzed with kymograph generation, using Kymo ToolBox ImageJ plugin. Colocalization was determined where a mRNA particle kymograph track overlapped with a track in the corresponding mitochondria kymograph. Moving particles were defined after movement of more than 10 µm in a specific direction during the movie 5-minute duration.

### Live imaging and analysis of N2a cells

N2a cells were seeded in 24-well glass-bottom plates at a density of 50,000 cells/well and allowed to grow for 24 hours. Cells were then transfected with MCP-YFP plasmid and one of the MS2 plasmids (0.25μg of each). Twenty-four hours after transfection, mitochondria were stained with 100 nM MitoTracker Red CMXRos (ThermoFisher) for 30 minutes at 37°C and washed with PBS before confocal live imaging. Images were captured using the LSM 710 inverted confocal microscope (Zeiss) with 63X NA 1.4 oil immersion objective lens. Z-stack images were acquired with 0.5μm z-step size and image pixel size of 0.1μm.

mRNA signal detection and mitochondria colocalization quantification were performed using Imaris. Mitochondria signal was masked and segmented using the built-in surface algorithm in Imaris. mRNA spots of 0.5 μm diameter were detected using the built-in spot detecting function. To take into account high signal variability between cells and to ensure correct signal detection, thresholds were chosen manually. Spot was considered colocalized if at least half of it overlapped with the mitochondria surface detected. Image processing for display proposes was done using FIJI. To filter background noise the MCP channel was subjected to the FTT bandpass filter and background subtraction. The mitochondrial signal channel was subjected to background subtraction.

### Statistical analysis

Statistical analysis was performed using build in tools in Microsoft Excel or GraphPad Prism, with specific tests indicated in each Figure legend.

## Notes

### Competing Interest Statement

The authors have declared no competing interest.

### Summary of Updates

Figure 1 was revised by inclusion of smFISH in motor neurons (panels K and L). Figure 3 was revised by inclusion of cellular fractionation and western blots (panels C and D), that were previously in the supplementary material. Text changes relevant to these additions (in Material and Methods and in Figure legends) were added. Supplementary material was removed (some was added to the main text). Affiliation of one of the authors was updated.

